# The neuropeptide Drosulfakinin regulates social isolation-induced aggression in *Drosophila*

**DOI:** 10.1101/646232

**Authors:** Pavan Agrawal, Damian Kao, Phuong Chung, Loren L. Looger

## Abstract

Social isolation strongly modulates behavior across the animal kingdom. We utilized the fruit fly *Drosophila melanogaster* to study social isolation-driven changes in animal behavior and gene expression in the brain. RNA-seq identified several head-expressed genes strongly responding to social isolation or enrichment. Of particular interest, social isolation downregulated expression of the gene encoding the neuropeptide *Drosulfakinin* (*Dsk*), the homologue of vertebrate cholecystokinin (CCK), which is critical for many mammalian social behaviors. *Dsk* knockdown significantly increased social isolation-induced aggression. Genetic activation or silencing of *Dsk* neurons each similarly increased isolation-driven aggression. Our results suggest a U-shaped dependence of social isolation-induced aggressive behavior on *Dsk* signaling, similar to the actions of many neuromodulators in other contexts.

**Data availability:** The raw sequence data from RNA-seq experiments has been deposited into the Sequence Read Archive (https://www.ncbi.nlm.nih.gov/sra) with accession number: PRJNA481582. Supplementary files and figures accompany this article.

## INTRODUCTION

Social isolation is a passive stressor that profoundly influences the behavior of social animals (Grippo et al., 2007; Hall et al., 1998; Wallace et al., 2009). Social isolation increases aggression in humans (Ferguson et al., 2005), rodents (Luciano and Lore, 1975; Ma et al., 2011; Wallace et al., 2009), and fruit flies (Hoffmann, 1989; Wang et al., 2008).

*Drosophila melanogaster* has been a successful model system for identifying the neural substrates of aggressive behavior (Asahina, 2017; Baier et al., 2002; Chen et al., 2002; Kravitz and Huber, 2003). Several conserved neuromodulators have been identified as key players in regulating aggression, including biogenic amines such as dopamine (Alekseyenko et al., 2013; Kayser et al., 2015), octopamine (Certel et al., 2007; Hoyer et al., 2008; Kayser et al., 2015; Williams et al., 2014; Zhou et al., 2008), and serotonin (Alekseyenko et al., 2010, 2014; Dierick and Greenspan, 2007); and neuropeptides including neuropeptide F (NPF; Asahina et al., 2014; Dierick and Greenspan, 2007) and tachykinin (Asahina et al., 2014). The associated receptors (Asahina et al., 2014) and neural circuits have been identified in some cases (Koganezawa et al., 2016).

Flies display aggression in a variety of settings, including male-male competition for females, territorial disputes, etc. (Asahina, 2017; Dow and Schilcher, 1975; Hoffmann, 1987; Jacobs, 1960; Kravitz and Fernandez, 2015). In this study, we sought to elucidate the circuit and genetic underpinnings of male aggression induced by deprivation of social interactions. Using an RNA-seq screen, we identified several candidate genes, most notably the neuropeptide Drosulfakinin (Dsk) (Chen and Ganetzky, 2012; Chen et al., 2012; Nichols et al., 1988; Söderberg et al., 2012), the homologue of the vertebrate cholecystokinin (CCK). CCK is well documented as a critical modulator of anxiety and aggression in a number of settings (Katsouni et al., 2013; Li et al., 2007; Panksepp et al., 2004; Vasar et al., 1993; Zwanzger et al., 2012). Dsk has been reported to modulate aggression in *Drosophila* (Williams et al., 2014), but many mechanistic details are lacking.

Here we use modulation of group size and isolation duration, RNA-seq, RNA interference (RNAi), and genetic activation or silencing of target neural populations to further elucidate the involvement of Dsk in aggressive behavior.

## RESULTS

### Social isolation induces transcriptional changes in male *Drosophila* heads

To probe the molecular mechanisms involved in regulating social isolation-induced behaviors, we performed RNA-seq on male flies that were housed either individually (single-housed, SH) or in groups of 20 (group-housed, GH) for four days in vials containing standard fly food. Flies were flash-frozen, whole heads isolated, RNA prepared and sequenced (N=2 biological replicates) (Figure 1A). Both SH and GH flies showed strong inter-replicate concordance: r = 0.964 and 0.965, respectively (Supplementary Figure S1A,B). Commonly used RNA-seq analysis methods utilize diverse models for dispersion, normalization and differentially-expressed gene (DEG) calling. To increase stringency of our DEG calling, we utilized three separate techniques, DEseq2 (Love et al., 2014), edgeR (Robinson et al., 2010), and EBseq (Leng et al., 2013), and compared their results (**Material and Methods**). We focused on genes identified by all three methods, which we considered to be robust hits.

**Figure 1.**
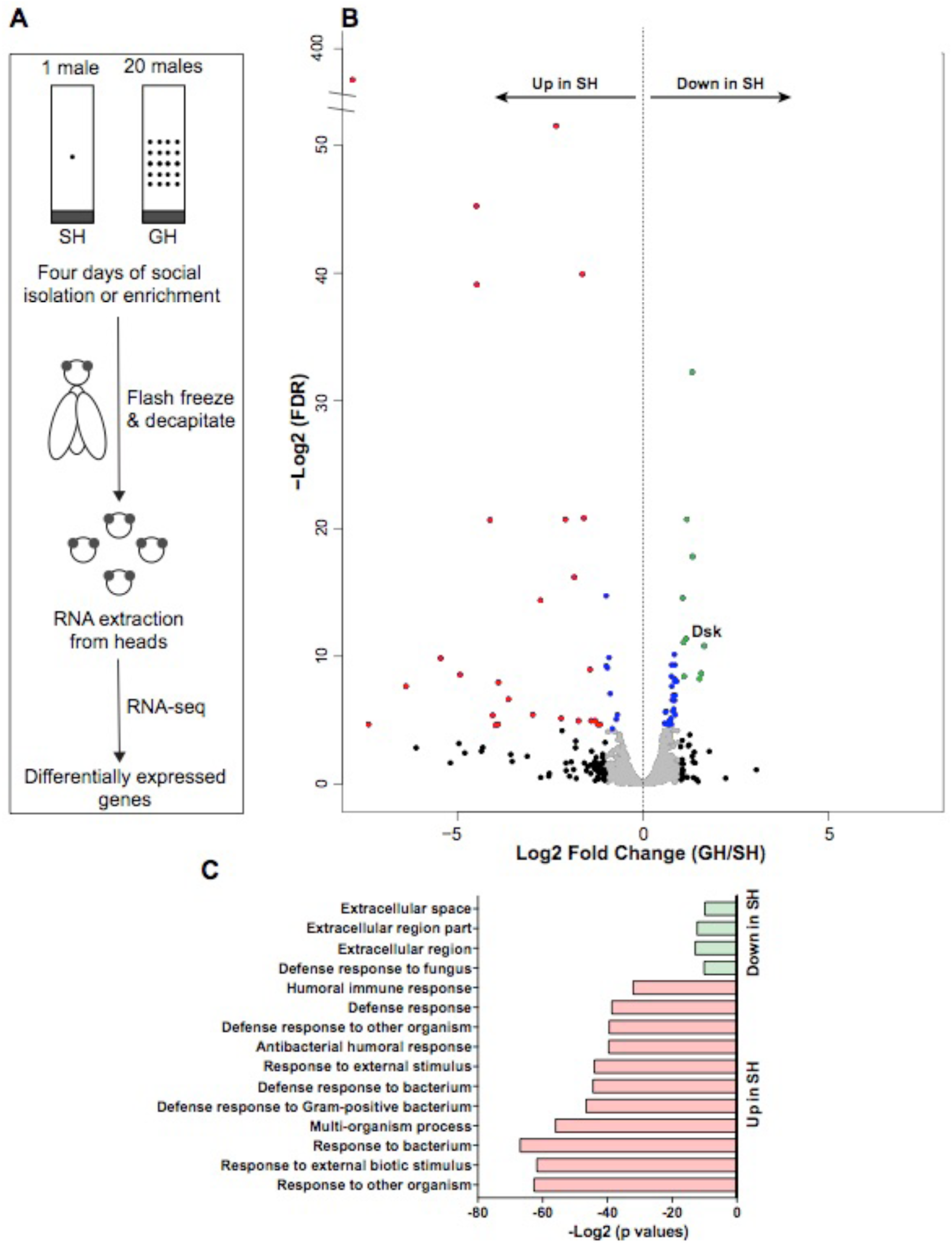
Transcriptional differences in male heads after single housing (SH) or group housing (GH). (A) Outline of the experimental paradigm. (B) Volcano plot of RNA-seq profile using DEseq2 for individual genes. Gray: FC<2; black: FC>2; blue: P<0.05; green: P <0.05 and +FC>2, red: P <0.05 and -FC>2; FC = fold change. (C) Enriched Gene Ontology categories for differentially-expressed genes obtained from DEseq2 analysis.

Using stringent criteria for differential expression, *i.e.* fold-change ≥ 2 and false discovery rate ≤ 0.05 (Figure 1B, Supplementary Table S1), 90 genes were selected by at least one method, and 25 by all three (Table 1). Most DEGs were related to the immune response (Table 1, Supplementary Tables S1, S2), which is consistent with the observation that social isolation leads to immune upregulation across the animal kingdom (Cole et al., 2015; Ellen et al., 2004; Powell et al., 2013). Many of these immune-related genes are commonly seen as DEGs in fly microarray and RNA-seq experiments (Carney, 2007; Ellis and Carney, 2011; Mohorianu et al., 2017; Wang et al., 2008) (Supplementary Table S3). We compared 90 social isolation-induced DEGs in whole fly heads with similar data generated specifically from FACS-purified dopaminergic neurons (Agrawal et al., 2019). Depending on the particular method used to identify DEGs, we found between 3 to 9 genes common between these two datasets, suggesting that social isolation regulates somewhat different set of genes in whole heads relative to dopaminergic neurons (Supplementary Table S3). Along these lines, perturbation of neural activity stimulates expression of distinct sets of genes in whole brain *versus* dopaminergic neurons (Chen et al., 2016).

**Table 1.** Genes identified as differentially expressed upon social isolation by three different RNA-seq analysis methods. The last column shows gene function from Flybase, or predicted from Pfam (http://pfam.xfam.org) classification or annotation of homologues in other species if not annotated in Flybase. A complete list of genes and GO analysis is included in Supplementary Tables S1 and S2.

| Genes down-regulated in response to social isolation |  |  |  |  |  |
| --- | --- | --- | --- | --- | --- |
| Flybase Gene ID | Gene symbol | DESeq2 | edgeR | EBseq | Annotation/known/predicted function |
| FBgn0039685 | Obp99b | Down | Down | Down | sensory perception of chemical stimulus |
| FBgn0000500 | Dsk | Down | Down | Down | neuropeptide |
| FBgn0032615 | CG6012 | Down | Down | Down | hydroxysteroid dehydrogenase |
| FBgn0036589 | CG13067 | Down | Down | Down | cuticle protein |
| FBgn0030398 | Cpr11B | Down | Down | Down | cuticle protein |
| FBgn0032507 | CG9377 | Down | Down | Down | epithelial protease |
| FBgn0040629 | CG18673 | Down | Down | Down | carbonic anhydrase 2 |
| FBgn0052282 | Drsl4 | Down | Down | Down | antifungal peptide |
| FBgn0035434 | Drsl5 | Down | Down | Down | antifungal peptide |
| Genes up-regulated in response to social isolation |  |  |  |  |  |
| FBgn0000278 | CecB | Up | Up | Up | antibacterial humoral response |
| FBgn0005660 | Ets21C | Up | Up | Up | defense response to bacterium |
| FBgn0032638 | SPH93 | Up | Up | Up | defense response to Gram-positive bacterium |
| FBgn0032639 | CG18563 | Up | Up | Up | epithelial protease |
| FBgn0265577 | CR44404 | Up | Up | Up | long non-coding RNA |
| FBgn0014865 | Mtk | Up | Up | Up | antifungal peptide |
| FBgn0036995 | Atk | Up | Up | Up | sensory cilium structural protein |
| FBgn0038299 | Spn88Eb | Up | Up | Up | fungal-induced protease inhibitor |
| FBgn0052185 | Edin | Up | Up | Up | defense response to Gram-negative bacteria |
| FBgn0037396 | CG11459 | Up | Up | Up | cathepsin-like protease |
| FBgn0010381 | Drs | Up | Up | Up | antifungal peptide |
| FBgn0043578 | PGRP-SB1 | Up | Up | Up | innate immune response |
| FBgn0010388 | Dro | Up | Up | Up | antibacterial peptide |
| FBgn0039593 | Sid | Up | Up | Up | response to bacterium |
| FBgn0262794 | CG43175 | Up | Up | Up | unannotated secreted peptide |
| FBgn0044812 | TotC | Up | Up | Up | response to bacterium |

In our data, very few (7 down-regulated, 4 up-regulated) not-obviously-immune transcripts were identified by all three RNA-seq analysis methods as being significantly modulated by social experience (Table1, Supplementary Table S1). Examples of genes down-regulated in SH males include the male-specific odorant-binding protein *Obp99b* (Anholt et al., 2003), hydroxysteroid dehydrogenase *CG6012*, and the neuropeptide *Drosulfakinin* (*Dsk*) (Figure 1B, Table1). Genes upregulated in SH males included the sensory cilium structural protein *Artichoke* (*Atk*), cathepsin-like protease *CG11459*, long non-coding RNA *CR44404*, and secreted peptide *CG43175* (Table1). Several of these transcripts have been identified in previous studies (Supplementary Table S3), albeit under different conditions (*e.g.* courtship, social defeat) (Barajas-Azpeleta et al., 2018; Carney, 2007; Ellis and Carney, 2011; Mohorianu et al., 2017; Wang et al., 2008) or by different techniques (*e.g.* microarray, RNA-seq). Of these stringently selected transcripts, only *Dsk* is central nervous system-specific and was selected for further study. *Dsk* expression differences were validated with qPCR on head-extracted RNA isolated from SH and GH males (Supplementary Figure S2A).

### *Dsk* knock-down affects social isolation-induced aggression

As CCK is known to regulate aggression, anxiety, and social-defeat responses in rodents (Katsouni et al., 2013; Li et al., 2007; Panksepp et al., 2004; Vasar et al., 1993; Zwanzger et al., 2012), we next tested specific phenotypic effects of *Dsk* modulation. Dsk localizes to the pars intercerebralis, parts of the protocerebrum and the sub-esophageal ganglion (Nichols, 1992; Nichols and Lim, 1996; Söderberg et al., 2012). In the pars intercerebralis, Dsk is expressed in the insulin-like peptide Dilp2-producing neurons (Söderberg et al., 2012). Faithful GAL4 driver lines are available for both *Dsk* (Asahina et al., 2014) and *Dilp2* (Rulifson, 2002) (Figure 2A, B). To quantify aggression, we counted lunges using the software package CADABRA (Dankert et al., 2009). Lunging, *i.e.* a fly rearing on its hind legs and snapping downward on its opponent, is a prominent aggressive behavior in *Drosophila* males (Hoffmann, 1987; Hoyer et al., 2008; Nilsen et al., 2004). Knockdown of *Dsk* in *Dsk-GAL4* neurons using RNA interference (RNAi) significantly increased lunges in SH flies relative to controls without RNAi insert (Figure 2C). Similar effects were observed upon *Dsk* knock-down using the *Dilp2-GAL4* driver (Figure 2E). Successful knockdown using pan-neuronal *elav-*GAL4 *^c155^* was confirmed by qRT-PCR (Supplementary Figure S2B). Thus, lowering *Dsk* expression in the pars intercerebralis increased aggressive lunging following social isolation.

**Figure 2.**
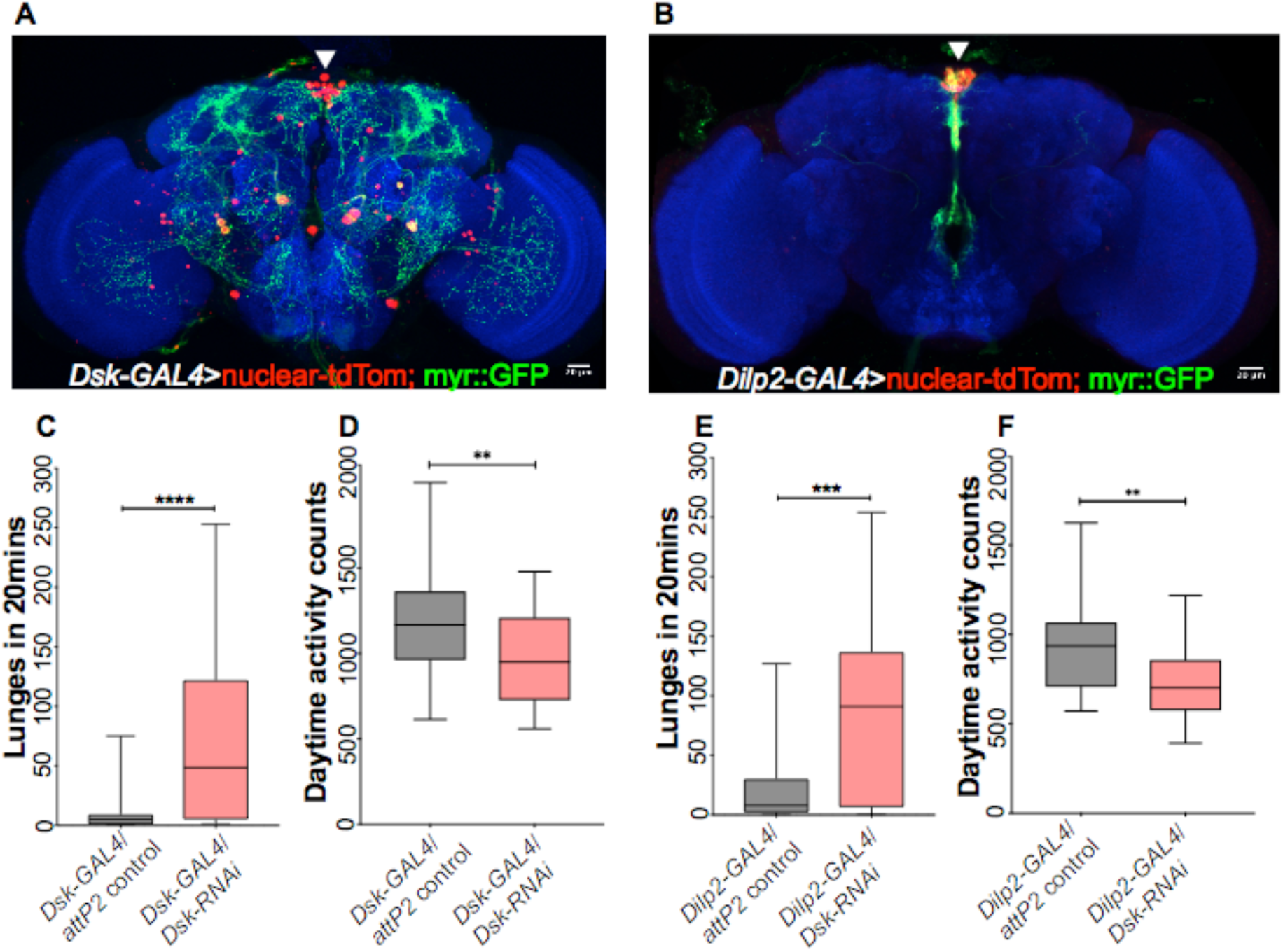
*Dsk* knockdown increases social isolation-induced aggression independently of overall activity levels. *Dsk* levels were reduced by driving the expression of *UAS-Dsk*-RNAi with *Dsk-GAL4* or *Dilp2-GAL4*. These drivers overlap in the pars intercerebralis region (white arrowhead, A and B). RNAi-mediated *Dsk* knockdown increased aggression in SH flies relative to the *attP2* background controls without RNAi insert (C, *Dsk-GAL4*, P<0.0001, N=38; E, *Dilp2-GAL4*, P=0.0004, N=26; Mann-Whitney U-test). *Dsk* knockdown significantly reduced the overall daytime activity of SH males (D, *Dsk-GAL4*, P=0.004, N=32; F, *Dilp2-GAL4*, P=0.002, N=32; Student’s t-test).

It was previously suggested that aggressive behaviors should be normalized to overall locomotor activity (Hoyer et al., 2008). Isolated wild-type flies sleep less and show greater levels of overall daytime activity (Ganguly-Fitzgerald et al., 2006) than GH flies. In contrast, isolated *Dsk*-knockdown flies show significantly reduced overall daytime activity (Figure 2D, F; for *Dsk-GAL4* and *Dilp2-GAL4*, respectively), with no effect in GH flies (Supplementary Figure S3). Thus, the observed increase in aggression in SH males upon *Dsk* knockdown arises despite decreased overall activity.

In *Drosophila*, Dsk has two receptors, CCK-like receptor (CCKLR)-17D1 and CCKLR-17D3 (Kubiak et al., 2002). Signaling through CCKLR-17D1, but not CCKLR-17D3, is responsible for larval neuromuscular junction growth and muscle contraction (Chen and Ganetzky, 2012; Chen et al., 2012). Since *GAL4* driver reagents are not available for several neuropeptide receptors, we used a MiMiC reporter line (Nagarkar-Jaiswal et al., 2015) available for *CCKLR-17D1* (**Material and Methods**) to ascertain its expression in the brain. We found reporter expression in several brains regions including in the pars intercerebralis (PI), which overlapped with *Dilp2* expression (Supplementary Figure S4A-C, A’-C’). Successful knockdown for both *CCKLR-17D1* and *CCKLR-17D3* was confirmed using *elav-*GAL4 *^c155^* by qRT-PCR (Supplementary Figure S4D). However, in *Dsk-GAL4* or *Dilp2-*GAL4 flies, knockdown of *CCKLR-17D1*, but not *CCKLR-17D3*, increased aggression of SH flies (Supplementary Figure S4E, F). This is consistent with results for the ligand *Dsk* and suggests that signaling of Dsk through its receptor CCKLR-17D1 in PI increases isolation-driven aggression.

### Social isolation is essential for *Dsk*-mediated aggression

To more precisely determine the interaction between social isolation and *Dsk*, we varied group size and isolation length from 1-20 flies and from 1-4 days, respectively. The presence of even a single other fly almost eliminated *Dsk* knockdown-evoked aggression, and aggression remained suppressed as group size increased (Figure 3 A, B). As few as 1-2 days of isolation modestly but significantly increased aggression in SH males in which *Dsk* was knocked down (Supplementary Figure S5A, B); the effect increased for up to 4 days.

**Figure 3.**
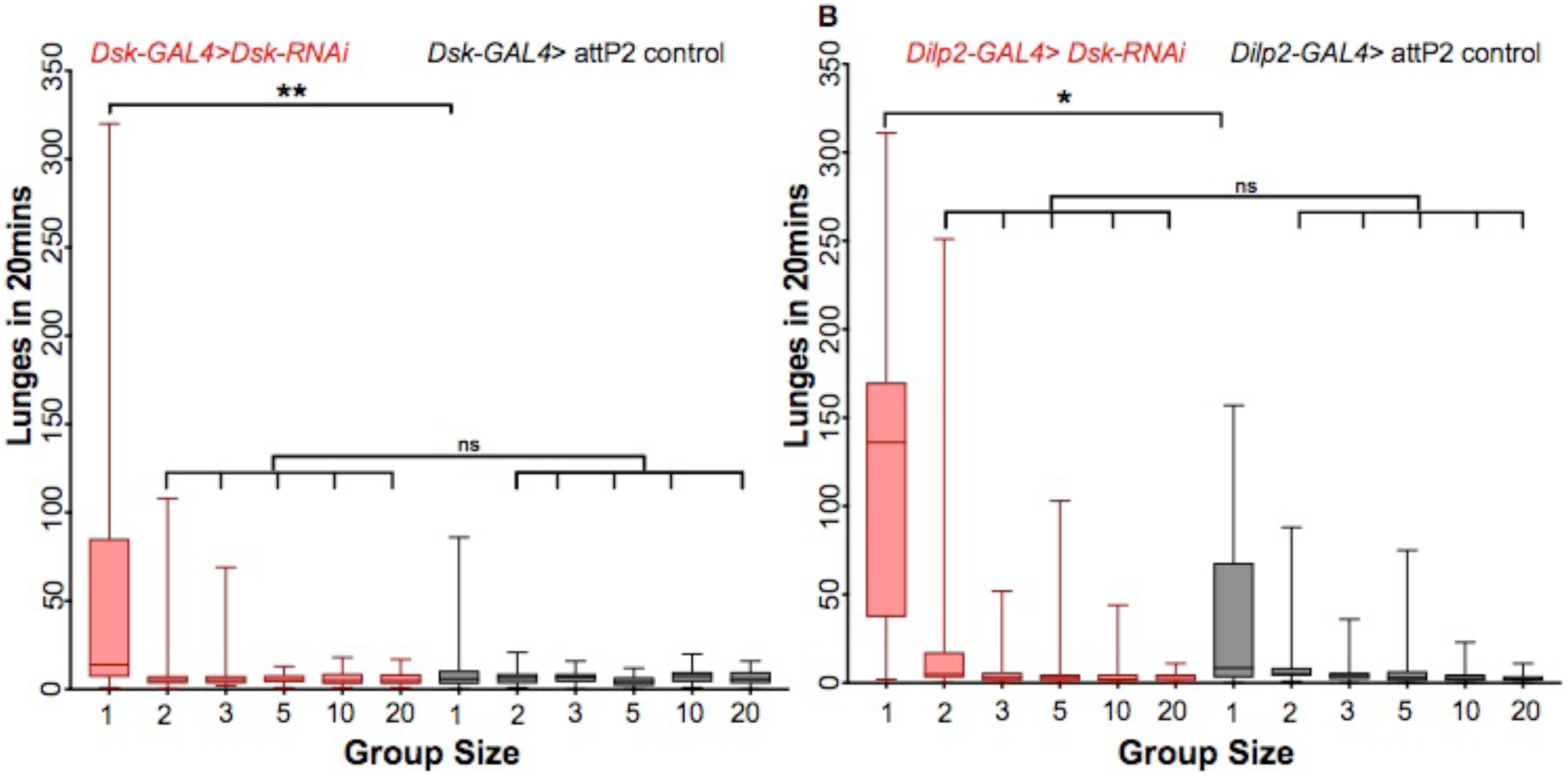
Social isolation is essential for aggression mediated by *Dsk* knockdown. (A) *Dsk-GAL4*/*Dsk-RNAi vs. Dsk-GAL4*/*attP2*-control, group size 1, **: P=0.0037. N=33-36. (B) *Dilp2-GAL4*/*Dsk-RNAi vs. Dilp2-GAL4*/*attP2*-control, group size 1, *: P=0.013. N=24-36. Presence of other males drastically reduced aggression as seen in group size 2 or greater. Kruskal-Wallis ANOVA with Dunn’s multiple comparison test.

### Both activation and silencing of *Dsk* neurons increase aggression

Having established the contribution of *Dsk* and its receptor *CCKLR-17D1* to aggression in SH males, we next explored the function of the neurosecretory cells themselves. Silencing of *Dsk* neurons with the inward rectifying potassium channel Kir2.1 (Baines et al., 2001) significantly increased lunging (Figure 4A), which is consistent with the involvement of *Dsk* and *CCKLR-17D1* signaling for promoting aggression in SH males. Surprisingly, genetic activation of *Dsk* neurons with the bacterial sodium channel NaChBac (Nitabach, 2006) also increased aggression (Figure 4B). GH flies showed very few lunges in all cases, indicating that social isolation is critical for aggression in our assays (Figure 4 A, B).

**Figure 4.**
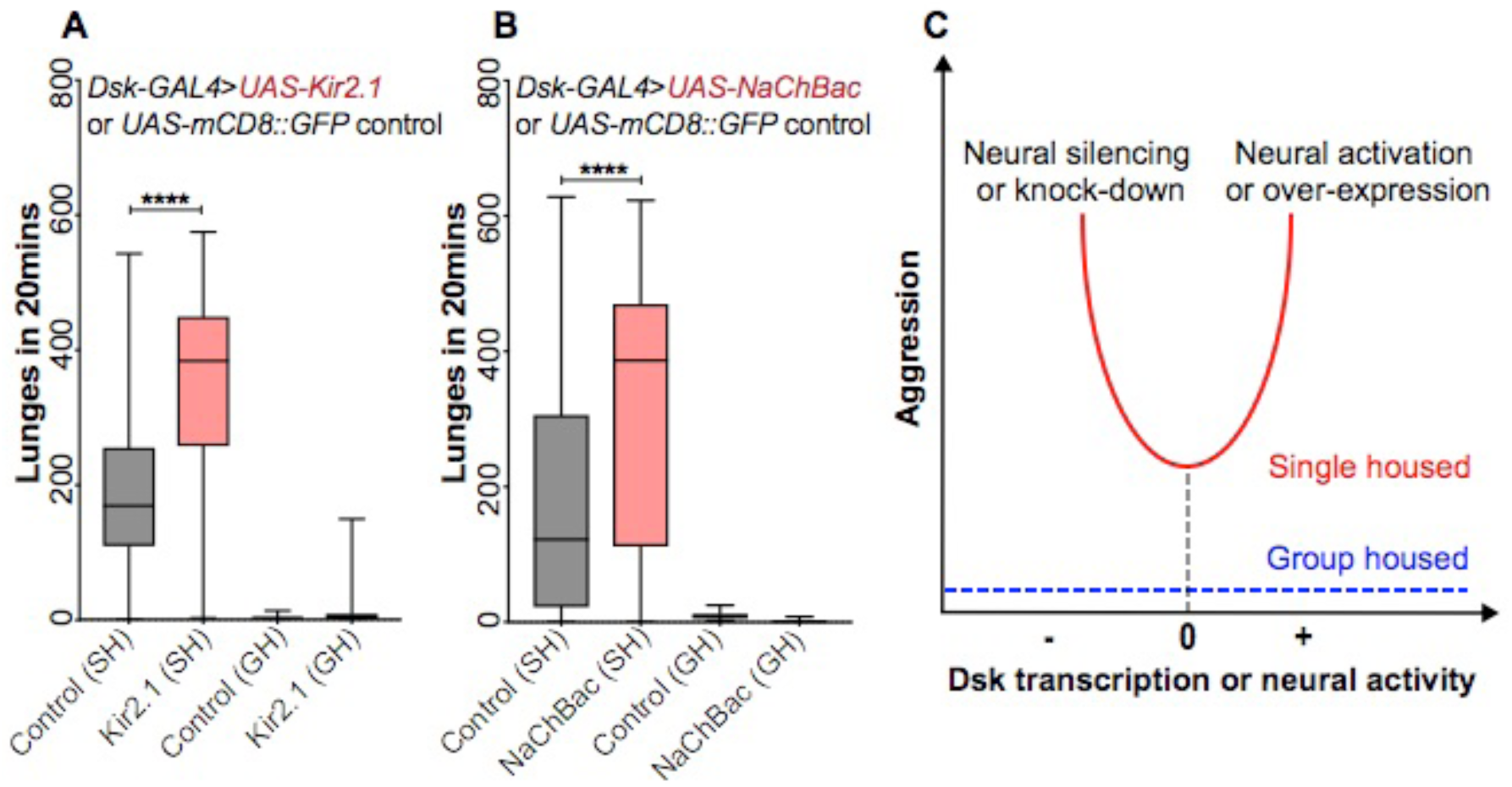
Genetic activation and silencing of *Dsk* neurons both increase aggression. (A) Silencing of *Dsk* neurons with *UAS-Kir2.1-EGFP* by *Dsk-GAL4* driver increases aggression in SH males (P< 0.0001, Mann-Whitney U-test, N=48). (B) Activation of *Dsk* neurons with *UAS-NaChBac* by *Dsk-GAL4* driver also increases aggression in SH males (P<0.0001, Mann-Whitney U-test, N=52). Controls in both (A) and (B) are *UAS-mCD8::GFP* transgene driven by *Dsk-GAL4.* (C) Schematic of putative U-shaped effect of *Dsk* neuron activity and transcript levels on aggression.

## DISCUSSION

We have shown that knockdown of the neuropeptide *Dsk* or its receptor *CCKLR-17D1* in the pars intercerebralis (PI) increases social isolation-driven aggression of male flies. Moreover, *Dsk* appears to act in a U-shaped fashion (Figure 4C), with both knockdown (our results) and overexpression (Williams et al., 2014) increasing aggression. We also showed that *Dsk* neuronal activity follows a similar trend, with both activation and silencing increasing aggression. This suggests that the primary role of these neurons in this context is indeed production and secretion of Dsk (Söderberg et al., 2012). Transcription factors in the fly PI neurons regulating aggression were recently identified (Davis et al., 2014), and it was shown that activation of PI neurons increases aggression. However, the downstream neuropeptides were not known (Thomas et al., 2015). Our findings identify *Dsk* as a key neuropeptide expressed in the PI region that regulates aggression. Further work will be required to delineate the aggression-modulating functions, if any, of other neuropeptides also secreted from the PI region.

A recent neural activation screen (Asahina et al., 2014) explored the role of neuropeptides in aggression in *Drosophila*, but investigated only group-housed flies. Intriguingly, Asahina *et al*. identified tachykinin signaling in the lateral protocerebrum and explicitly ruled out involvement of *Dsk* neurons (Asahina et al., 2014) in GH fly aggression. Thus, male-male aggression in GH and SH flies appears to be controlled by different neuropeptides in different brain regions. The absence of *Dsk* neurons from the screen results in GH flies (Asahina et al., 2014), combined with our results showing suppressed aggression in GH flies regardless of *Dsk* transcription or neural activity, suggests a mechanism that overrides *Dsk* function.

Downregulation of Dsk receptor *CCKLR-17D1* in *Dsk/Dilp2* neurons also increased aggression, consistent with the observation that some neuropeptidergic neurons, *e.g.* those for neuropeptide F (Shao et al., 2017; Wen et al., 2005), neuropeptide Y (Qi et al., 2016) and FMRFamide (Ravi et al., 2018) have receptors to modulate their signaling in an autocrine manner. However, pan-neuronal down-regulation of *CCKLR-17D1* receptor did not affect aggression (data not shown), suggesting potential antagonistic effects outside *Dsk/Dilp2* neurons.

### Hormetic regulation of behaviors

The U-shaped (“hormetic”) response of the aggression phenotype to both Dsk levels and *Dsk*^+^ neuronal activity are similar to such responses seen for NPF (Asahina et al., 2014; Dierick and Greenspan, 2007) and dopamine (Alekseyenko et al., 2013) neurons in *Drosophila* aggression. Such effects are not unexpected, given the ubiquity of such hormetic responses in neuromodulator signaling pathways (Baldi and Bucherelli, 2005; F.Flood et al., 1987; Monte-Silva et al., 2009) and receptors in general (Calabrese and Baldwin, 2001). At the level of individual G-protein coupled receptors, such U-shaped responses (low-dose agonism, high-dose antagonism) arise directly from equations considering receptor expression level and the effects of receptor activation on downstream signaling pathways (Kohn and Melnick, 2002). At the circuit level, it is thought that such U-shaped responses help to maintain neuronal activity patterns, and the resulting behaviors, near homeostatic optima, with deviations resulting in negative feedback (Arnsten et al., 2012; Brunel and Wang, 2001; Herman, 2013).

### Social experience modulates gene expression in *Drosophila* heads

There have been a number of prior studies on the genetic basis of aggression in *Drosophila*, many of them performed with DNA microarrays – which record counts for specific transcripts of interest – rather than with RNA-seq, which counts all transcripts within cells. Four such studies have been performed in recent years, each identifying a large number of putative aggression-related genes: (Edwards et al., 2006) found 1672 such transcripts, (Dierick and Greenspan, 2006) found 149, (Wang et al., 2008) found 183, and (Tauber, 2010) found 339. It should be noted that these four studies used very different experimental methods: (Edwards et al., 2006) and (Dierick and Greenspan, 2006) bred flies for aggressive behavior over several generations, and isolated mRNA for microarray analysis at a time unrelated to aggression events; thus, these genes are generally high in the selected flies. (Wang et al., 2008) analyzed single-housed and group-housed flies, again irrespective of specific aggression events. (Tauber, 2010), meanwhile, isolated pairs of aggressive flies and obtained mRNA for sequencing directly following bouts, looking for aggression bout-driven gene expression. Given the substantial differences in experimental design, and the imperfect reliability of microarray quantification, it is perhaps unsurprising that of the 1672, 149, 183, and 339 differentially expressed genes in each study, respectively, there were only 2 in common to all four studies: the olfactory binding protein Obp99b and CG13794, an unannotated transporter. Obp99b appeared amongst our 25 most significant hits, whereas CG13794 did not appear to be differentially expressed at all in our assays. Given that the involvement of Dsk in aggression is quite context-specific, for instance Asahina *et al*. explicitly ruled out of involvement in aggression of group-housed flies (Asahina et al., 2014), it is perhaps unsurprising that it was not found in several of the screens. In fact, the only one of these four studies to uncover Dsk was the one that utilized socially isolated flies (Wang et al., 2008), strengthening the notion that Dsk specifically links social isolation to aggression. It was this link with social behavior that drew our attention to Dsk, and indeed our experiments bear out that this function is mediated through activity in the central brain. The pars intercerebralis has been shown to be the seat of regulation of many other social and sexually dimorphic behaviors (Belgacem and Martin, 2002; Luo et al., 2014; Mattaliano et al., 2007; Terhzaz et al., 2007).

At the other end of the spectrum, olfactory inputs are ubiquitous and odor processing through olfactory receptors factors into essentially every fly action. Along with Dsk, Obp99b was the most down-regulated gene in our single-housed males (Table 1, Supplementary Table S1). Obp99b was also picked up in two studies of other social behaviors: courtship-exposed males (Carney, 2007) and male competition for mates (Mohorianu et al., 2017). Olfactory binding proteins (OBPs) are secreted by support cells in the antennal trichoid sensilla to assist in odorant binding and recognition by olfactory receptors (Galindo and Smith, 2001; Larter et al., 2016). A critical role for Obp76a (Lush) in recognition of the pheromone cis-vaccenyl acetate (cVA) by olfactory receptor Or67d, and in driving aggression following acute pheromone exposure, has been established (Billeter and Levine, 2015; Wang and Anderson, 2010). However, following chronic cVA exposure, the pheromone also activates a second receptor, Or65a, which then inhibits Or67d glomeruli and decreases aggression (Liu et al., 2011). The OBP mediating cVA recognition by Or65a is currently unknown, but appears not to be Lush (Laughlin et al., 2008). It is possible that the OBP identified in all the screens discussed, *i.e.* Obp99b, recognizes cVA for signaling through Or65a; it is also possible that it recognizes other odorants. Given its ubiquity in screens for social behaviors, we speculate that the molecules recognized are likely pheromones. Obp99b is one of the most male-specific transcripts identified (Fujii and Amrein, 2002), and indeed it had been previously discovered as a gene in the sex-determination cascade, under its previous name Turn on Sex-Specificity (Tsx) (Wolfner, 2003). In this set of experiments, we selected Dsk for investigation because of its link to social context; however, the precise function of Obp99b warrants closer study, given its probable role in pheromone detection.

Other prominent hits from our screen appear interesting, as well (Table 1, Supplementary Table S1). The third most isolation-driven down-regulated transcript, *CG6012*, appears to encode a hydroxysteroid dehydrogenase. Such enzymes have been shown to be critical for pheromone production in insects, with the related enzyme CG1444 (Spidey) processing both ecdysone and related cuticular hydrocarbons (Chiang et al., 2016). The Turandot peptides, annotated as stress-response genes, also appear to play sex-specific roles in behaviors such as courtship. Turandot A and Turandot C, both up-regulated in isolated males in our study (Supplementary Table S1), are greatly female-enriched, and Turandot C, in particular, is up-regulated in female flies following playing of an attractive, conspecific courtship song over a speaker (Immonen and Ritchie, 2012). Finally, we would note that new roles have recently been proposed for transcripts annotated as encoding antimicrobial peptides (*e.g.* Diptericin B, DptB), specifically the modulation of long-term memory (Barajas-Azpeleta et al., 2018). It is possible that some of the transcripts annotated as antimicrobial peptides are instead (or in addition) memory regulators with roles in social behaviors. Indeed, playing synthetic attractive, conspecific *versus* aversive, heterospecific courtship song dramatically lowered expression of the ostensibly antimicrobial peptides Attacin-A, Attacin-C, DptB, Drosocin, and Immune-induced molecule 18 (Immonen and Ritchie, 2012). Intriguingly, all of these molecules were increased in males following aversive social isolation in our study (Figure 1 B, C, Supplementary Table S1). The only other molecule shared between the two studies, Methuselah-like 8, showed the opposite pattern: up-regulated in females hearing attractive song and down-regulated in males after aversive isolation. Of course, given the wealth of bacteria, fungi, viruses and other microbes present in and on flies and their food, it is probable that many annotated antimicrobial peptides are indeed responding to differences in pathogen load composition between SH and GH flies. But the observation that many putatively antimicrobial molecules respond strongly to stimuli (*e.g.* synthetic courtship song) that do not involve alteration of their physical environment in any way indicates that these molecules have more sophisticated functions in the brain, and may encode valence (attractive, aversive) of social interactions.

### Dsk and its homologue CCK have evolutionarily conserved role in regulating aggression

In mammals, the Dsk homologue cholecystokinin (CCK) and its receptors regulate aggression, anxiety, and social-defeat responses (Katsouni et al., 2013; Li et al., 2007; Panksepp et al., 2004; Vasar et al., 1993; Zwanzger et al., 2012). For instance, intravenous injection of the smallest isoform, CCK-4, in humans reliably induces panic attacks (Bradwejn et al., 1991; Tõru et al., 2010) and is often used to screen anxiolytic drug candidates. However, in other contexts, such as in mating (Bloch et al., 1987) and juvenile play (Burgdorf et al., 2006), CCK encodes strong positive valence. CCK colocalizes with dopamine in the ventral striatum, and microinjection of CCK into the rat nucleus accumbens phenocopies the effects of dopamine agonists, increasing attention and reward-related behaviors (Vaccarino, 1994), further supporting its role in positive valence encoding. CCK actions differ across brain regions, in a context-dependent manner. For instance, time pinned (negative-valence) during rough-and-tumble play correlated with increased CCK levels in the posterior cortex and decreased levels in hypothalamus (Burgdorf et al., 2006). However, lower hypothalamic CCK also correlated with positive-valence play aspects including dorsal contacts and 50 kHz ultrasonic vocalizations. Thus, CCK can encode both positive- and negative-valence aspects of complex behaviors differentially across the brain. As with many neuromodulators (Calabrese, 2001; Cools and D’Esposito, 2011; Joëls, 2006), CCK appears to act in a U-shaped fashion, with increases and decreases of signaling from baseline levels often producing similar phenotypes (Burgdorf et al., 2006; Calabrese and Baldwin, 2003; Ding and Bayer, 1993; Kõks et al., 1999; Kulkosky et al., 1976).

Taken together, our results suggest an evolutionarily-conserved role for neuropeptide signaling through the drosulfakinin pathway (homologue of cholecystokinin) in promoting aggression. Intriguingly, this pathway only seems active in socially-isolated flies; in socially-enriched flies, aggression is controlled by tachykinin (*a.k.a.* Substance P) signaling. The PI region, in which the *Dsk*/*Dilp2* neurons reside, has considerable similarities with the hypothalamus (Hartenstein, 2006), a brain region crucial for regulating aggression in mammals (Gregg and Siegel, 2001; Haller, 2013; Kruk et al., 1984; Lin et al., 2011; Lipp and Hunsperger, 1978; Toth et al., 2010), with the most relevant activity localized to the ventrolateral subdivision of the ventromedial hypothalamus (Lin et al., 2011), where CCK neurons reside (Fulwiler and Saper, 1985). Thus, the predominant aggression-regulating mechanism in rodents bears strong homology to the fly pathway regulating aggression of socially-deprived, but not socially-enriched, individuals.

## MATERIALS AND METHODS

### Fly stocks & rearing

Flies were reared on standard food at 25°C and 65% relative humidity with a 12-h light/dark cycle. For behavioral and molecular experiments, flies were collected within 24-48 hours of eclosion and group housed (GH) or single housed (SH) for four days, unless mentioned otherwise. The following fly strains were obtained from the Bloomington stock center: *Dsk-*GAL4 (#BL51981; Asahina et al., 2014); *Dilp2-*GAL4 (*Dilp2 a.k.a. Ilp2*; #BL37516); *elav-*GAL4*^c155^* (#BL458). For *Dsk-* and *Dilp2-*GAL4 expression analysis a fluorescent reporter (Etheredge et al., 2018) carrying *pJFRC105-10XUAS-IVS-NLS-tdTomato* in VK00040 (a gift of Barret D. Pfeiffer, Rubin lab, Janelia Research Campus) and *pJFRC29-10XUAS-IVS-myr::GFP-p10* in attP40 (Pfeiffer et al., 2012) was used. For examining CCKLR-17D1 expression a MiMIC reporter (Nagarkar-Jaiswal et al., 2015) (#BL 61771; y[1] w[*] Mi{PT-GFSTF.2}CCKLR-17D1[MI03679-GFSTF.2]) was used. For aggression and qPCR assays, comparisons were made between equivalent genetic backgrounds. The stock used for neural silencing *pUAS-Kir2.1-EGFP* and its corresponding control *pJFRC2-10XUAS-IVS-mCD8::GFP* were obtained from the Fly Facility Shared Resource at Janelia Research Campus. *Dsk-GAL4*, *Dilp2-GAL4* and *elav*-*GAL4^c155^* and stocks used for neural activation (*UAS-NaChBac*, #BL9466 and control *UAS-mCD8::GFP*, #BL5130) were outcrossed for 6-7 generations into *w*; Berlin background. The following Transgenic RNAi Project (TRiP) RNAi lines (Perkins et al., 2015) were obtained from the Bloomington stock center: *Dsk*-RNAi (#BL25869); *CCKLR-17D1*-RNAi (#BL27494); *CCKLR-17D3*-RNAi (#BL28333) and the *attP2* background control without RNAi insert (#BL36303). To negate effects of the mini-white gene on aggression (Hoyer et al., 2008), male progenies containing (*w* y[1] v[1]) were obtained by crossing virgin females of various GAL4 drivers and males of TRiP RNAi for *Dsk*, *CCKLR-17D1* and *CCKLR-17D3* or corresponding background controls.

### Immunohistochemistry and imaging

Fly brains were dissected in cold 1X PBS and fixed in 2% paraformaldehyde (in 1X PBS) at room temperature for one hour on a Nutator, washed 4 times for 20 min each in PAT (1X PBS, 0.5% PBS Triton, 1% BSA) at room temperature, blocked for one hour at room temperature with blocking buffer (PAT + 3% Normal Goat Serum) and incubated with primary antibodies, diluted in blocking buffer, overnight on a Nutator at 4°C. The primary antibodies used were: mouse anti-GFP (Sigma-Aldrich, #G6539, 1:200 dilution); Rabbit anti-DsRed (Clontech, 632496, 1:500 dilution), rat anti-DN-cadherin (Developmental Studies Hybridoma Bank, DNEX#8, 1:50 dilution), mouse anti-Flag (Sigma-Aldrich, #F1804, 1:100 dilution); and Rat anti-Dilp2 (1:800 dilution, Eric Rulifson). This was followed by 4 washes for 20 min each in PAT, and incubation overnight on a Nutator at 4°C with secondary antibodies diluted in blocking buffer. The secondary antibodies were all from Molecular Probes and used at 1:500 dilution:

Alexa Fluor 488 anti-mouse (A11029), Alexa Fluor 568 anti-rabbit (A11036), Alexa Fluor 568 anti-rat (A11077) and Alexa Fluor 633 anti-rat (A21094). Brains were then washed 4 times for 20 min each in PAT at room temperature, 1 time for 20 min in 1X PBS and mounted with VECTASHIELD mounting medium (Vector Laboratories, H-1000). Samples were imaged on a Zeiss 800 confocal laser-scanning microscope.

### RNA extraction, library preparation and sequencing

Male Canton-S flies collected within 24-48 hours of eclosion were group housed (GH) or single housed (SH) for four days and flash-frozen during the afternoon and stored at −80°C until RNA extraction. 10-15 flies were vortex-decapitated and heads were collected on dry ice. Heads were lysed in Trizol, and total RNA was extracted using a Zymo Direct Zol kit (Zymo Research, USA: #R2051), in-tube DNAse digestion was performed using a Turbo DNA free kit (Thermo Fisher Scientific, USA: #AM1907), and RNA was purified using a Zymo RNA Clean and Concentrator kit (Zymo Research, USA: #R1013) as per manufacturer’s instructions. External RNA Controls Consortium (ERCC) spike-ins were added, and RNA was processed for sequencing using Ovation RNA-Seq System V2 (Nugen Technologies, USA: #7102-32) and Ovation Rapid DR Multiplex System 1-8 (Nugen Technologies, USA: #0319-32) as per manufacturer’s instructions. Two biological replicates were performed for each condition. Paired-end 100 bp sequencing reads were obtained using Illumina Hi-seq 2500 (Illumina, San Diego, CA).

### RNA-seq analysis

All reads were trimmed with Trimmomatic 0.36 at a minimum read length of 50 and average read quality across a sliding window of 15. Trimmed reads were mapped to *Drosophila* genome version r6.03 with STAR (Dobin et al., 2013) with default settings. Pairwise differential expression analysis was performed with DESeq2, EBseq, and edgeR following instructions given in the respective R package’s workflow. Genes that were differentially expressed at stricter than an adjusted (corrected for multiple testing using the Benjamini-Hochberg method) P-value of 0.05 and fold-change greater than 2 were used for further analysis. Gene Ontology (GO) analysis was performed on enriched genes using GOrilla (http://cbl-gorilla.cs.technion.ac.il/). ~5,000 genes expressed in fly heads were used as background for GO analysis and obtained from FlyBase (http://www.flybase.org). The raw data from RNA-seq experiments has been deposited into the Sequence Read Archive (https://www.ncbi.nlm.nih.gov/sra) with accession number: PRJNA481582.

### qPCR validation

RNA was extracted from heads of flies as described in previous sections. Genotype and age of flies used for qPCR was matched to their corresponding behavioral assay. After RNA extraction, cDNA was prepared using a SuperScript VILO Master Mix kit (Thermo-Fisher Scientific, USA: #11755050). qPCR was performed using Brilliant III Ultra-Fast QPCR Master Mix (Agilent Technologies: #600880) on a StepOne Plus Real-Time PCR System (Thermo-Fisher Scientific, USA: #4376600). The following hydrolysis probes (Applied Biosystems, Life Technologies, USA) were used: RPL32 (*Dm02151827_g1*: #4331182) as an endogenous control, *Dsk* (*Dm02152147_s1*: #4351372), CCKLR-17D1 (*Dm01813942_g1*: # 4448892) and CCKLR-17D3 (*Dm01813944_m1*: # 4448892) as test probes.

### Aggression assay

The assay was performed essentially as described before (Dankert et al., 2009; Kim et al., 2018). In brief, males of a given genotype were introduced as a pair by gentle aspiration into single wells (16 mm diameter and 10mm height) of 12-well chambers. Chamber floors were made from 2.25% w/v agarose in commercial apple juice and 2.5% (w/v) sucrose. Walls of the arena were covered with Fluon (BioQuip) to prevent flies from climbing the walls. Flies were allowed to acclimatize to the arena for 5 minutes, and then fights were recorded for 20 minutes at 30 frames per second. All fights were performed during the morning activity peak within 2.5 hours of lights on, at 25°C and ~40% relative humidity. Lunges were counted by CADABRA (Caltech Automated Drosophila Aggression-Courtship Behavioral Repertoire Analysis) software (Dankert et al., 2009).

### Locomotor activity analysis

Flies of various genotypes that were previously SH or GH for 4 days were anesthetized briefly by carbon dioxide and transferred into 5mm × 65mm transparent plastic tubes with standard cornmeal dextrose agar media. For recording locomotion levels, *Drosophila* activity monitors (Trikinetics, Waltham, USA) were kept in incubators at 25°C with 65% relative humidity on a 12-h light/dark cycle. Flies were allowed one night to acclimatize to the activity monitor, and then data was collected in 1-minute bins for 24 hours (day-time plus night-time activity) as described before (Donelson et al., 2012).

### Statistical analysis

Statistical analysis of behavioral data was performed using Prism 7 (Graph pad software). Aggression data is usually non-normally distributed and appropriate non-parametric tests were chosen. For activity data parametric test were chosen. Unless specified, we used ANOVA (non-parametric or parametric, as appropriate), followed by appropriate *post hoc* tests of significance. We used Mann-Whitney U (*a.k.a.* Wilcoxon rank-sum), Kruskal-Wallis and Student’s t-tests of significance, as appropriate.

## Supplementary Figures with legends

**Supplementary Figure S1.**
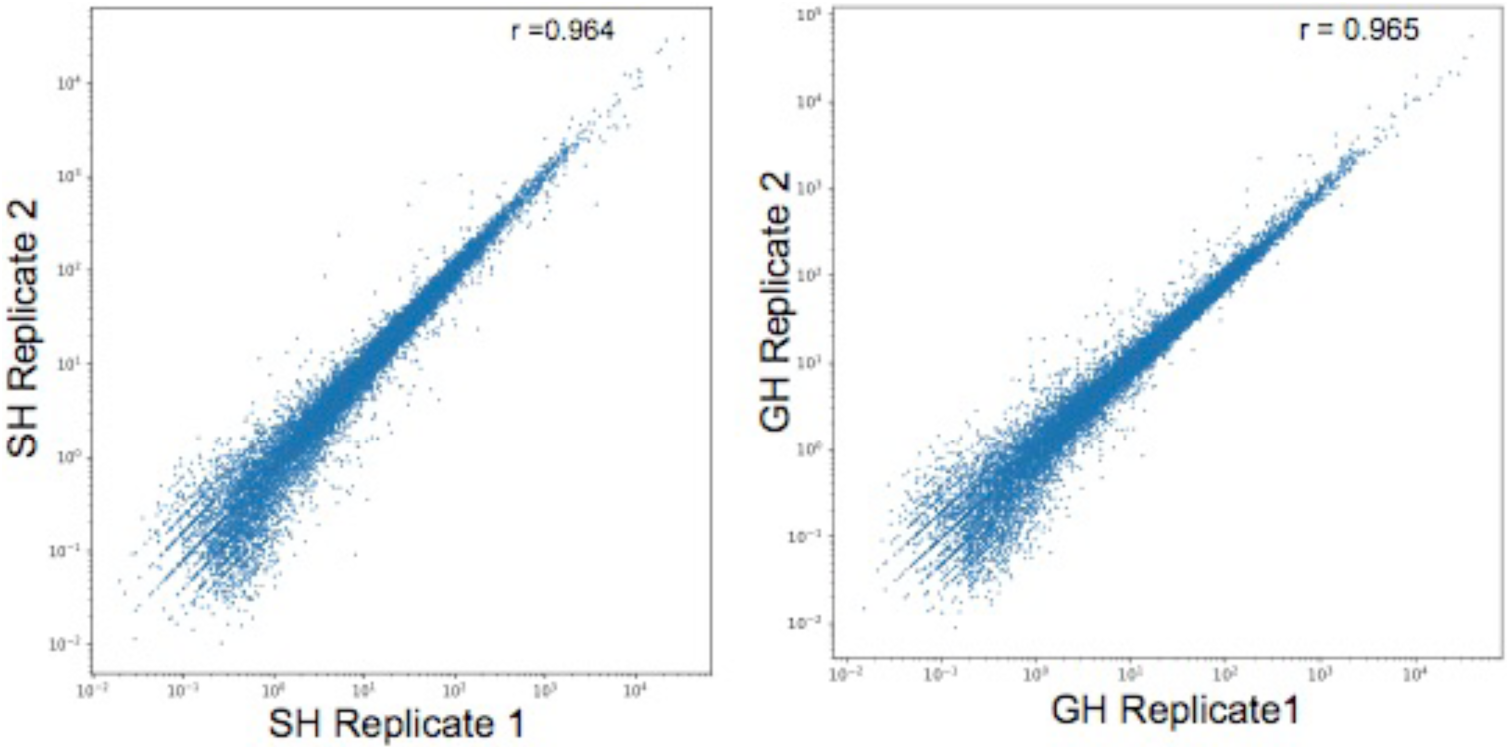
Replicate concordance of RNA-seq for GH and SH datasets. Pearson r-coefficients were calculated on the Transcripts Per Million (TPM) values for independent biological replicates (N=2) for RNA-seq data obtained from heads of (A) single-housed (SH) and (B) group-housed (GH) flies.

**Supplementary Figure S2.**
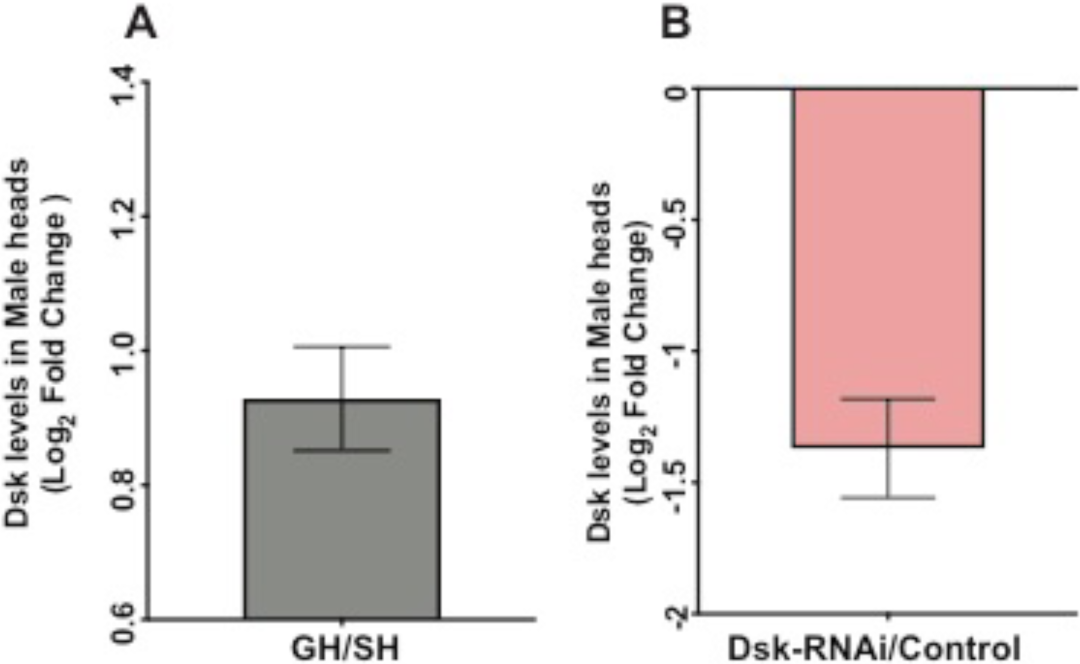
qPCR confirmation of Dsk upregulation upon group-housing and RNAi-mediated knockdown. (A) qPCR confirmation of *Dsk* transcriptional up-regulation in GH fly heads compared to SH fly heads. (N= 6 biological replicates). (B) qPCR confirmation of *Dsk* knockdown. *Dsk-RNAi* was driven pan-neuronally by *elav-GAL4^c155^* and compared with controls without RNAi insert driven by *elav-GAL4^c155^*. (N= 5 biological replicates). Y-axis shows Log_2_ Fold change of *Dsk* expression calculated using the ΔΔCt method; error bars show mean ± SEM. *Rpl32* was used as an endogenous control.

**Supplementary Figure S3.**
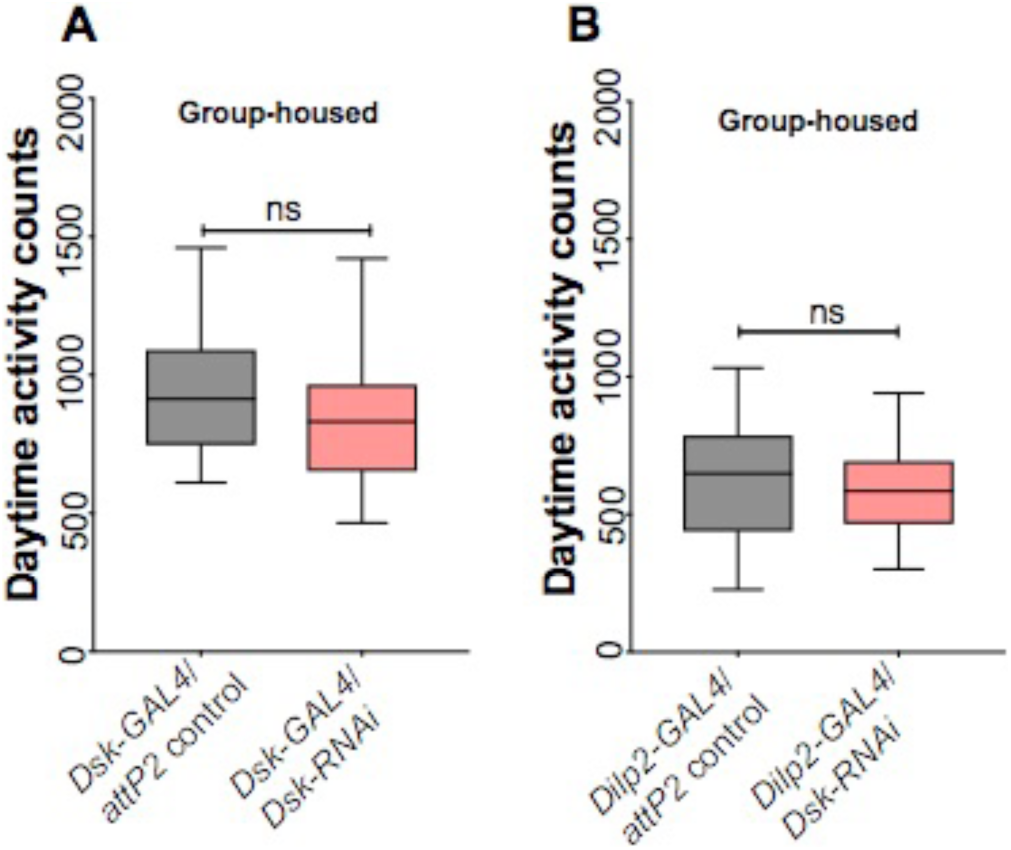
Group-housed flies do not show differences in overall daytime activity upon *Dsk* knockdown. Daytime activity is not significantly affected when *Dsk* was down-regulated using (A) *Dsk*-GAL4 driver GH, Student’s t-test; N=32; and (B) *Dilp2*-GAL4 driver GH, Student’s t-test; N=32.

**Supplementary Figure S4.**
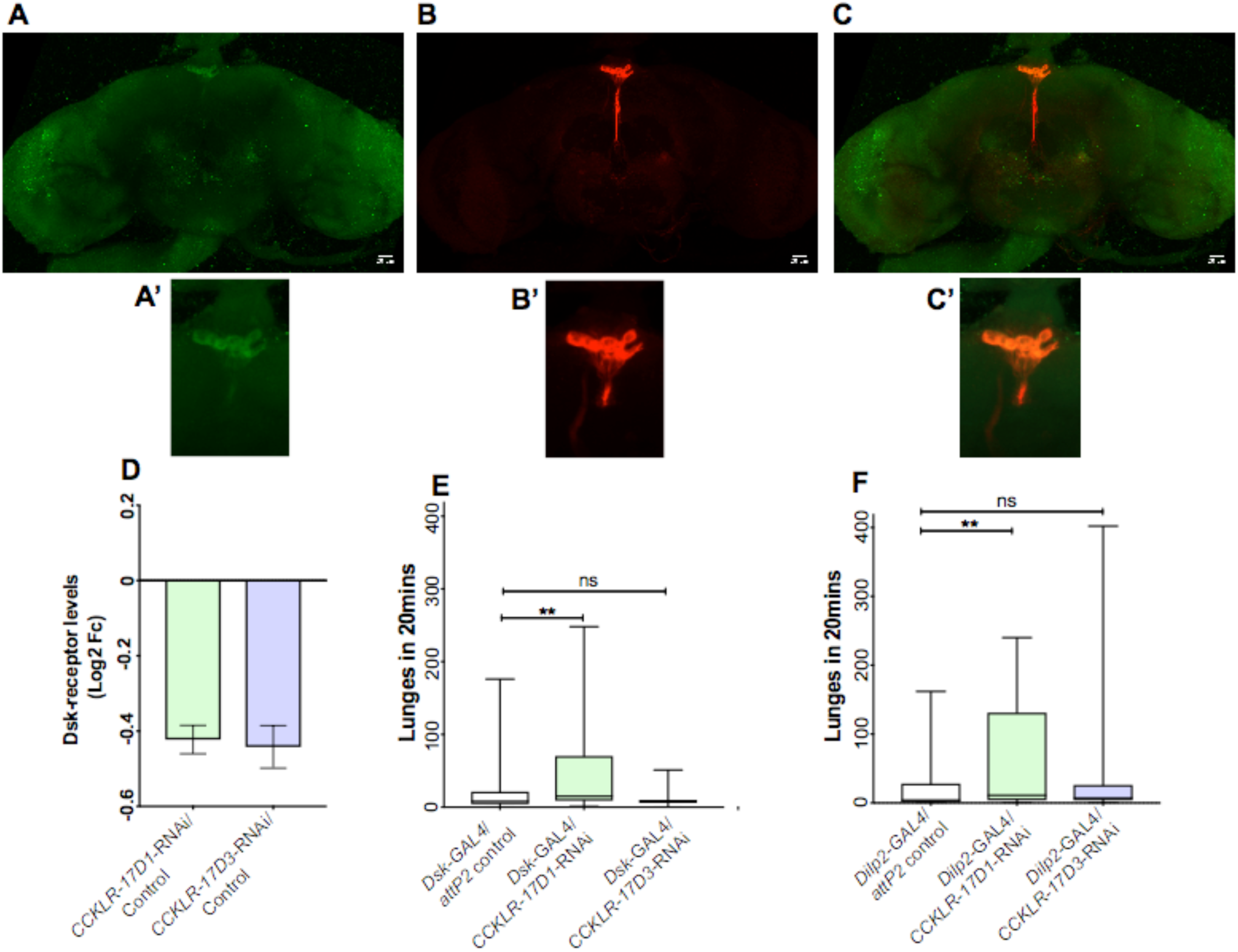
Aggression is mediated by the CCKLR1-17D1 receptor in peptidergic neurons. *CCKLR-17D1* expression in the brain ascertained using a MiMiC reporter line. (A) *CCKLR-17D1* reporter (green) is expressed in various brain regions including in PI region, where (B) *Dilp2* is expressed (red). (C) Overlap in PI region is shown. Scale bar is 20 μm. A’, B’ and C’ show zoomed images from PI region. (D) qPCR confirmation of *CCKLR-17D1* and *CCKLR-17D3* knockdown. RNAi constructs against *CCKLR* were driven pan-neuronally by *elav-GAL4^c155^* and compared with controls without RNAi insert driven by *elav-GAL4^c155^*(N=3 biological replicates). Y-axis shows Log_2_ Fold change of *Dsk* expression calculated using the ΔΔCt method; error bars show mean ± SEM. *Rpl32* probe was used as an endogenous control. (E,F) Knockdown of *CCKLR-17D1* led to increased aggression in both (E) *Dsk*-GAL4> *CCKLR-17D1*-RNAi (P= 0.003, Kruskal-Wallis ANOVA with Dunn’s multiple comparison test, N= 44-48) and (F) *Dilp2*-GAL4> *CCKLR-17D1*-RNAi (P= 0.0045, Kruskal-Wallis ANOVA with Dunn’s multiple comparison test, N=47-48). Aggression was not significantly affected when *CCKLR-17D3* was down-regulated in either GAL4 driver line.

**Supplementary Figure S5.**
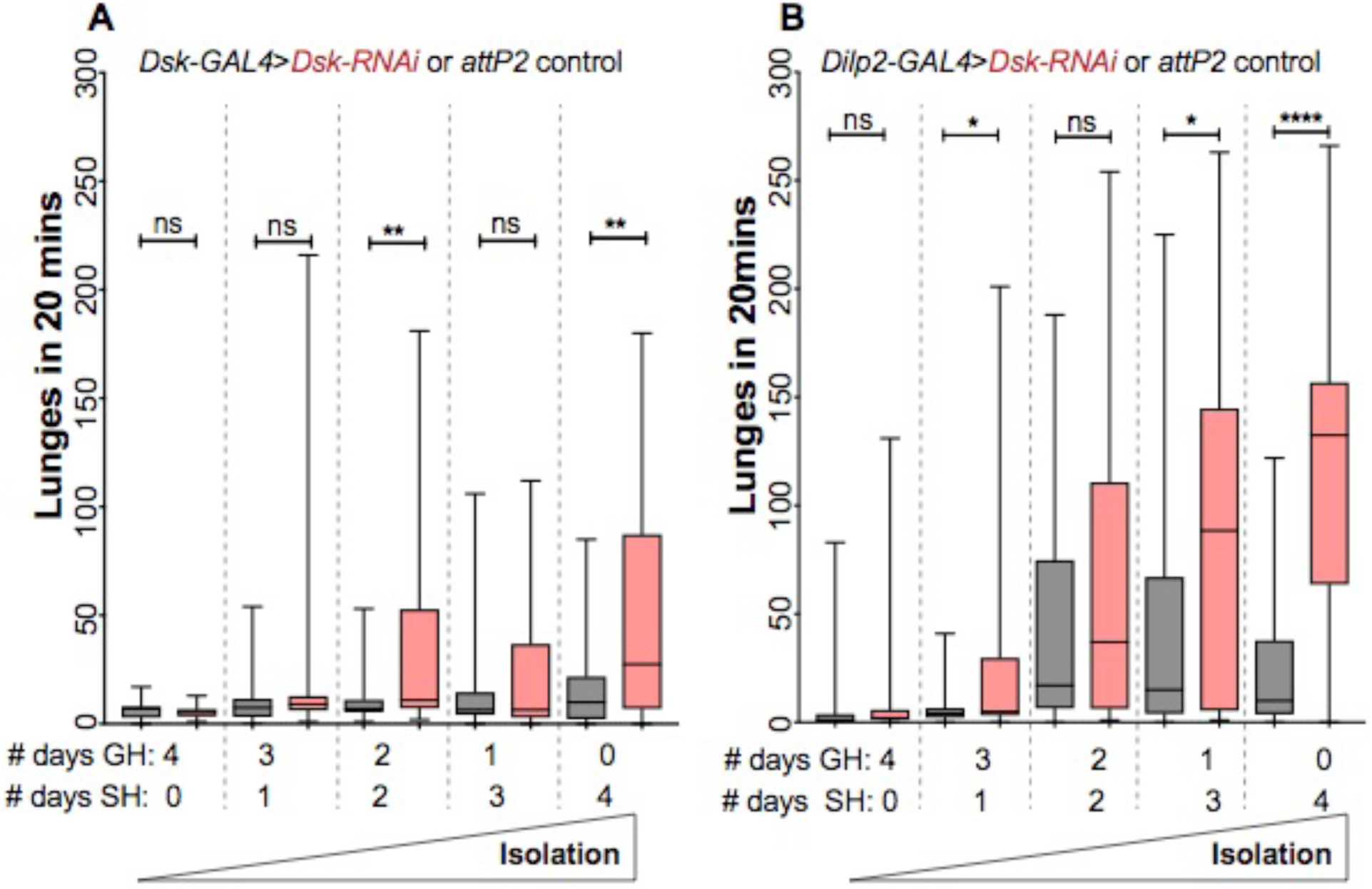
Effect of increased degree of social-isolation on *Dsk* knockdown-mediated aggression. Lunge numbers versus degree of isolation, GAL4 driver line, and *Dsk*-RNAi *vs*. *attP2* background control. Flies were group housed (20 males/vial) followed by single-housing. for varying degree. (A) *Dsk*-GAL4. **, P =0.008 for 2-days GH> 2-days SH; **; P = 0.0096 for 4 days of SH. N= 25-41. Mann-Whitney U-test. (B) *Dilp2*-GAL4. *; P = 0.043 for 3-days GH > 1-day SH; *; P = 0.013 for 1-day GH > 3-day SH and ****; P <0.0001 for 4 days of SH. N= 31-49. Mann-Whitney U-test.

## ACKNOWLEDGMENTS

We would like to thank Ulrike Heberlein for her support throughout this work, helpful discussions and critical reading of this manuscript. We thank Serge Picard, Andy Lemire and Janelia Quantitative Genomics for sequencing, Clement Kent and Anton Schulmann for helpful discussions related to RNA-seq analysis. We thank Karen Hibbard and Fly Facility Shared Resource at Janelia for help with fly genetics, Herman Dierick (Baylor College of Medicine, USA), Barret Pfeiffer, Gerald Rubin and Jack Etheredge (Janelia) for fly stocks. We also thank Lisha Shao, Mark Eddison and Jasper Simon (Janelia) for helpful discussions.

## AUTHOR CONTRIBUTIONS

PA conceptualized the project, designed and performed experiments, analyzed data and wrote the manuscript. DK performed RNA-seq analysis. PC performed immunostaining and imaging. LLL designed experiments, analyzed the data and wrote the manuscript. All authors reviewed the final MS.

